# Desialylated platelets promote hepatocyte proliferation via the ERK1/2 signaling pathway

**DOI:** 10.64898/2026.07.26.740293

**Authors:** Ippei Noboruo, Tomofumi Nakamura, Mayu Okumura, Tomoka Nishijima, Hiroki Inada, Yasuhito Tanaka, Tatsuya Kawaguchi, Masao Matsuoka, Jun-ichirou Yasunaga, Mitsuhiro Uchiba, Yukinori Kozuma

**Affiliations:** Department of Medical Technology, Kumamoto Health Science University, Kumamoto, Japan; Department of Hematology, Rheumatology, and Infectious Diseases, Graduate School of Medical Sciences, Kumamoto University, Kumamoto, Japan; Department of Gastroenterology and Hepatology, Faculty of Life Sciences, Kumamoto University, Kumamoto, Japan; Department of Blood Transfusion and Cell Therapy, Kumamoto University Hospital, Kumamoto, Japan

**Keywords:** Desialylated platelets, hepatocyte proliferation, HepG2 cells, hCLiPs, ERK1/2

## Abstract

Platelets are increasingly recognized as active regulators of tissue repair and liver regeneration beyond their classical roles in hemostasis and thrombosis. Loss of terminal sialic acid from platelet surface glycoproteins, a process known as desialylation, occurs during platelet aging or activation and has been linked to platelet clearance via the asialoglycoprotein receptor (ASGPR) on hepatocytes. However, the mechanisms by which desialylated platelets (D-plts) directly stimulate hepatocyte proliferation remain poorly understood. This study aimed to elucidate the proliferative effects of D-plts on hepatocytes and to identify the underlying signaling mechanisms. D-plts were generated and co-cultured with hepatocyte models exhibiting low or absent levels of asialoglycoprotein receptor 1 (ASGPR1) expression, including HepG2 cells, HuH-7 cells, and human chemically induced liver progenitors. Hepatocyte proliferation was assessed, and the roles of platelet-derived factors and downstream signaling pathways were investigated. Co-culture with D-plts significantly increased hepatocyte proliferation in all three cell models compared with the corresponding controls. Moreover, supernatants derived from stimulated D-plts also significantly enhanced hepatocyte proliferation, suggesting that soluble platelet-derived factors contribute to this effect. Mechanistically, the proliferative effects were mediated predominantly through the ERK1/2 signaling pathway rather than the JAK-STAT pathway in both hepatocytes co-cultured with D-plts and those treated with D-plt-derived supernatants. In conclusion, our findings demonstrate that D-plts directly promote hepatocyte proliferation through an ASGPR-independent pathway, in which ERK1/2 signaling plays a central role. These results highlight a novel mechanism through which platelet desialylation may contribute to liver regeneration.

**Graphical Abstract:**

(A) Desialylated platelets are readily activated and release increased amounts of EGF, promoting hepatocyte proliferation via the EGF–ERK signaling pathway.
(B) Normal platelets show lower reactivity and reduced EGF release than desialylated platelets, resulting in weaker hepatocyte proliferation.

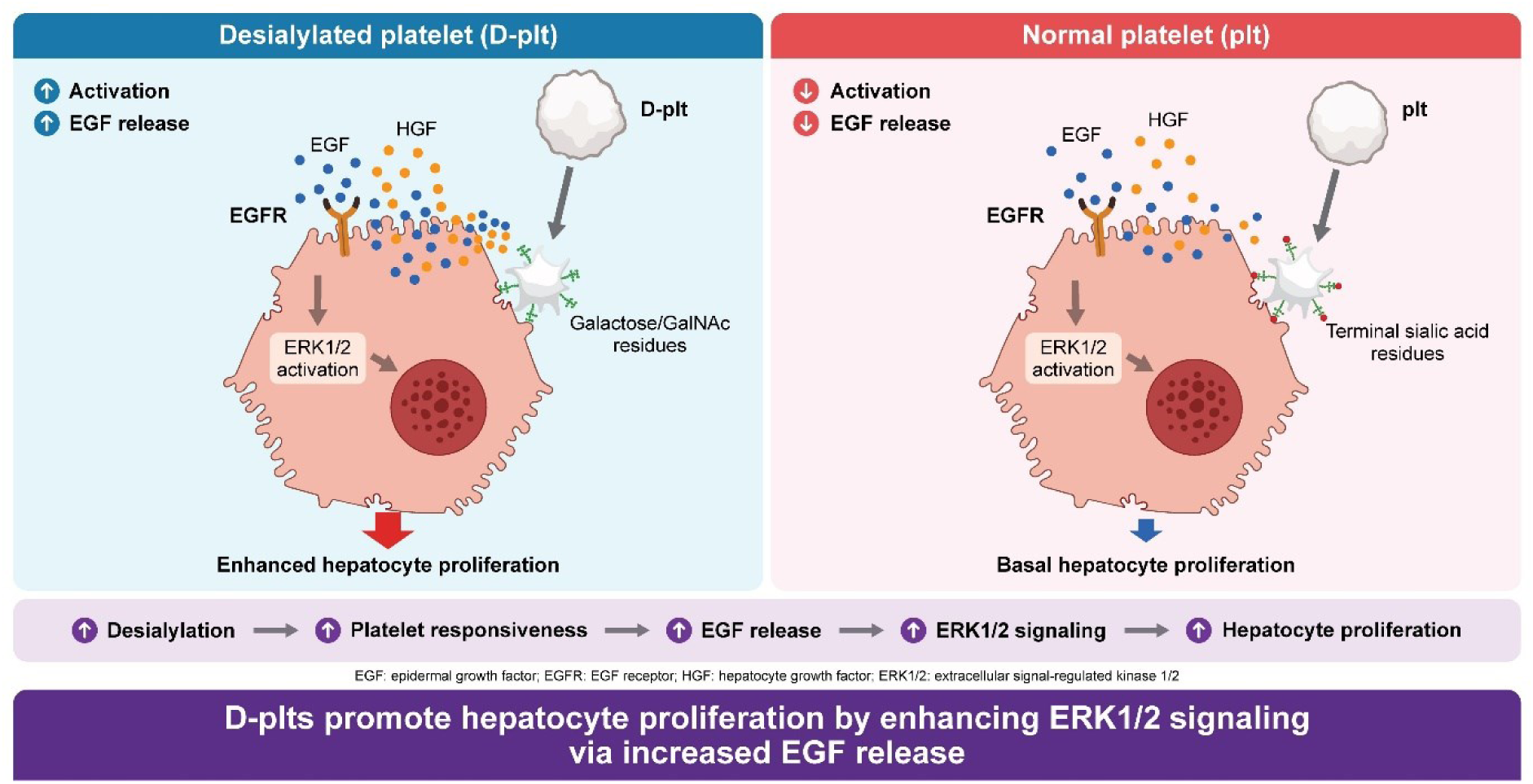

## Introduction

Platelets have long been recognized as primary mediators of hemostasis and thrombosis. However, accumulating evidence indicates that they also function as multifunctional effector cells with important roles in tissue repair and organ regeneration [1–5]. Among the key determinants of these platelet functions, the glycan structures at the terminal regions of platelet membrane glycoproteins have attracted considerable attention. In particular, terminal sialic acid plays a critical role in regulating platelet lifespan and clearance by serving as a recognition signal for hepatic receptors [6,7]. During aging or activation, platelets progressively lose terminal sialic acid from surface glycoproteins, a process known as desialylation [8]. This process has traditionally been regarded as a marker of platelet clearance; however, recent evidence suggests that desialylation may also contribute to a functional shift in platelets toward tissue homeostasis and regeneration [9].

Grozovsky *et al*. demonstrated that desialylated platelets (D-plts) are recognized and internalized by the Ashwell–Morell receptor (AMR), also known as the asialoglycoprotein receptor (ASGPR), on hepatocytes [10]. This interaction activates the Janus kinase 2 (JAK2)–signal transducer and activator of transcription 3 (STAT3) signaling pathway, inducing thrombopoietin (TPO) production and establishing a feedback loop for platelet homeostasis. In addition, Kurokawa *et al*. reported that platelets promote early liver regeneration in a mouse partial hepatectomy model [11]. However, whether platelet desialylation directly modulates hepatocyte proliferation and the underlying molecular mechanisms remains poorly understood.

The current model of TPO production is based on receptor-mediated endocytosis, and assumes sufficient ASGPR expression in healthy hepatocytes. In contrast, ASGPR expression is markedly downregulated under pathological conditions such as severe liver injury or hepatocellular carcinoma [12]. This raises the possibility that platelet-mediated signaling to hepatocytes may occur through mechanisms independent of ASGPR-mediated internalization.

Therefore, the present study aimed to investigate whether such a non-canonical pathway exists. To address this, we utilized HepG2 cells, HuH-7 cells, and human chemically induced liver progenitors (hCLiPs), which exhibit minimal or no functional expression of ASGPR. ASGPR is widely recognized as a marker of mature hepatocytes, and its expression is low in immature hepatocytes, such as hepatic progenitor cells, but increases during differentiation into mature hepatocytes [13]. Using this model system, we sought to elucidate the molecular mechanisms through which D-plts directly or indirectly promote hepatocyte proliferation.

## Results

### Generation and functional characterization of desialylated platelets from healthy donors

To generate experimental D-plts, we prepared washed platelets treated with neuraminidase (NEU), which removes sialic acid from platelet membrane glycoproteins (Fig 1A). The generated D-plts were characterized by flow cytometry (FCM) using *Ricinus communis* agglutinin-I (RCA-I) and glycoprotein (GP) Ibα (CD42b) antibodies. RCA-I recognizes terminal galactose residues exposed on GPIbα following desialylation. NEU treatment of platelets from healthy donors resulted in a marked increase in RCA-I–positive platelets, with more than 90% of platelets becoming RCA-I positive (Fig 1B). This finding indicates that NEU treatment efficiently generated D-plts. Importantly, NEU treatment did not alter the expression of CD42b, a platelet surface glycoprotein, and no change was observed in P-selectin expression, a marker of platelet activation (Figs 1C and D). These results indicate that NEU treatment specifically induces desialylation without affecting platelet integrity or activation status.

**Fig 1.**
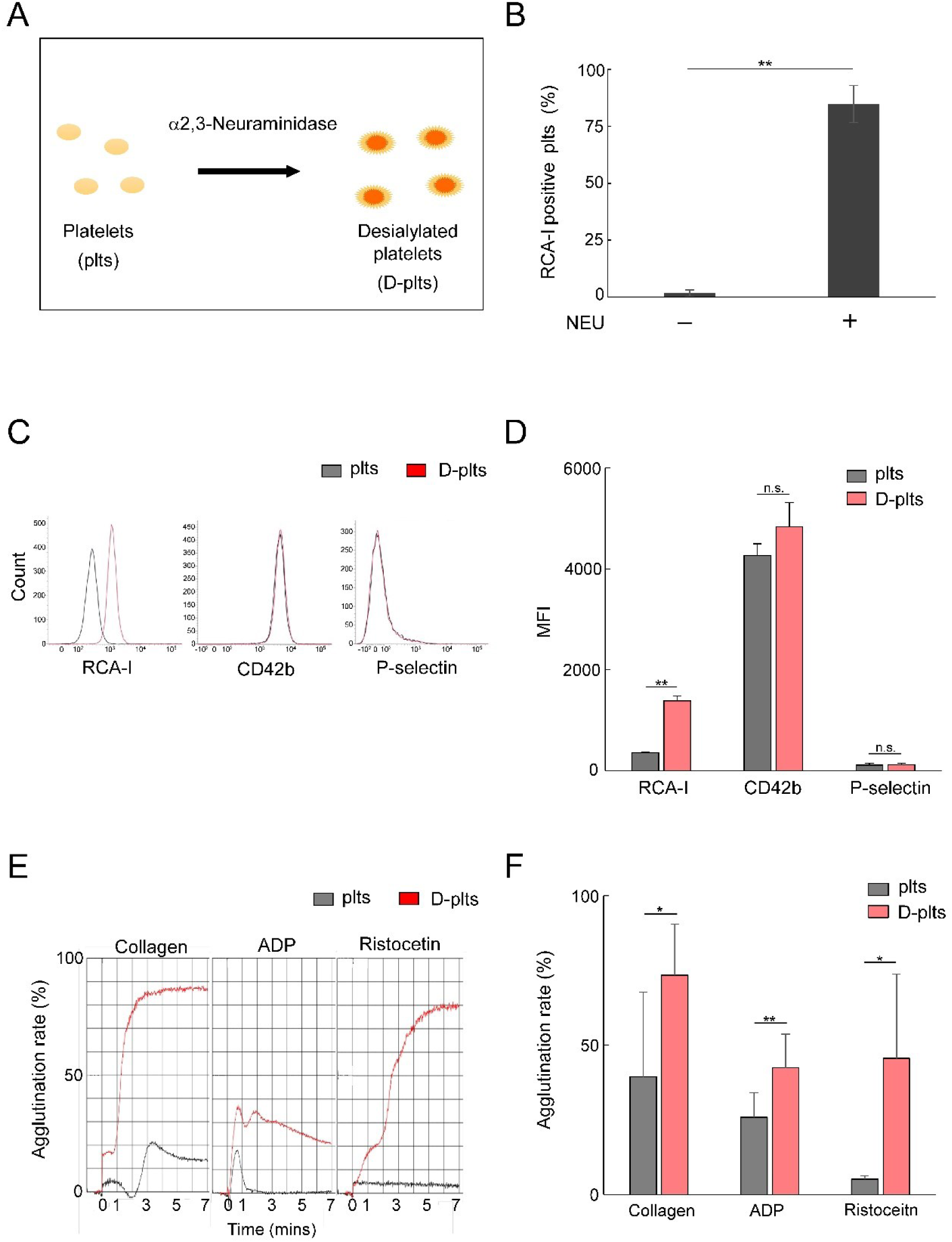
Generation and characterization of desialylated platelets. (**A**) Schematic representation of experimental desialylated platelet (D-plt) generation by treating washed platelets with or without neuraminidase (NEU) at 37°C for 15 min. **(B)** Platelets were stained with FITC-conjugated RCA-I lectin, and the proportion of RCA-I-positive platelets was quantified by flow cytometry (FCM). The percentage of D-plts was quantified in samples obtained from three healthy donors after treatment with or without NEU. (**C, D**) Changes in P-selectin (CD62P) and CD42b expression following NEU treatment. (**E, F**) Platelet aggregation following NEU treatment in response to collagen, ADP, and ristocetin. Data are presented as mean ± standard deviation (SD) (n = 3). Statistical comparisons between two groups were performed using the paired Student’s *t* test. Statistical significance was defined as \**p* < 0.05 and \*\**p* < 0.01.

To evaluate platelet function after NEU treatment, platelet aggregation assays were performed. Stimulation with collagen, adenosine diphosphate (ADP), and ristocetin resulted in enhanced platelet aggregation compared with the control group (Figs 1E and F). These findings suggest that D-plts are not fully activated but are more susceptible to activation.

### Changes in morphology and quality of stored platelets

To evaluate spontaneous changes in platelet quality during storage, we investigated platelets stored in blood bags. The platelet count decreased from day 3 of storage compared with day 0 (S1A Fig), and the pH progressively decreased from day 10 onward (S1B Fig). In contrast, the mean platelet volume (MPV) increased after 7 days (S1C Fig). The maximum aggregation rate was 62% on day 0 when platelets were stimulated with collagen plus ADP. Platelet aggregation gradually decreased on days 3, 7, and 14 of storage (S1D Fig). In addition, swirling patterns were attenuated after 10 days of storage and completely disappeared after 14 days (S1E Fig). Platelet aggregates within the storage bags were observed on days 10–14 (S1E Fig).

We also evaluated the morphological changes in D-plts using scanning electron microscopy (SEM). No apparent morphological differences were observed between untreated platelets and D-plts (S1F and G Fig). We further analyzed the occurrence of D-plts during storage by measuring the percentage of RCA-I–positive platelets using FCM (S2A–C Fig). The proportion of RCA-I–positive platelets increased significantly after 3 days (6.9 ± 0.6%) and 7 days (14.6 ± 0.7%) of storage compared with day 0 (3.0 ± 0.4%), suggesting that D-plts become detectable after 3 days of storage (S2A-C Fig). The expression of CD42b gradually decreased over the first 7 days of storage (S2D Fig). In contrast, membrane P-selectin expression, a marker of platelet activation, progressively increased on days 0, 1, 3, and 7, of storage (S2E Fig). These findings indicate that platelet-based experiments conducted within a 7-day storage period are unlikely to be compromised in terms of platelet function and quality.

### The effect of normal and desialylated platelets on HepG2 cell growth

ASGPR is a glycan-recognizing receptor expressed on hepatocytes that functions as a heterocomplex composed of ASGPR1 and ASGPR2 [14–16]. ASGPR1 expression was not detected on the surface of HepG2 cells (Fig 2A). When HepG2 cells were directly co-cultured with D-plts, cell growth was significantly greater than that in the control and normal platelets (plts) groups (Figs 2B–D).

**Fig 2.**
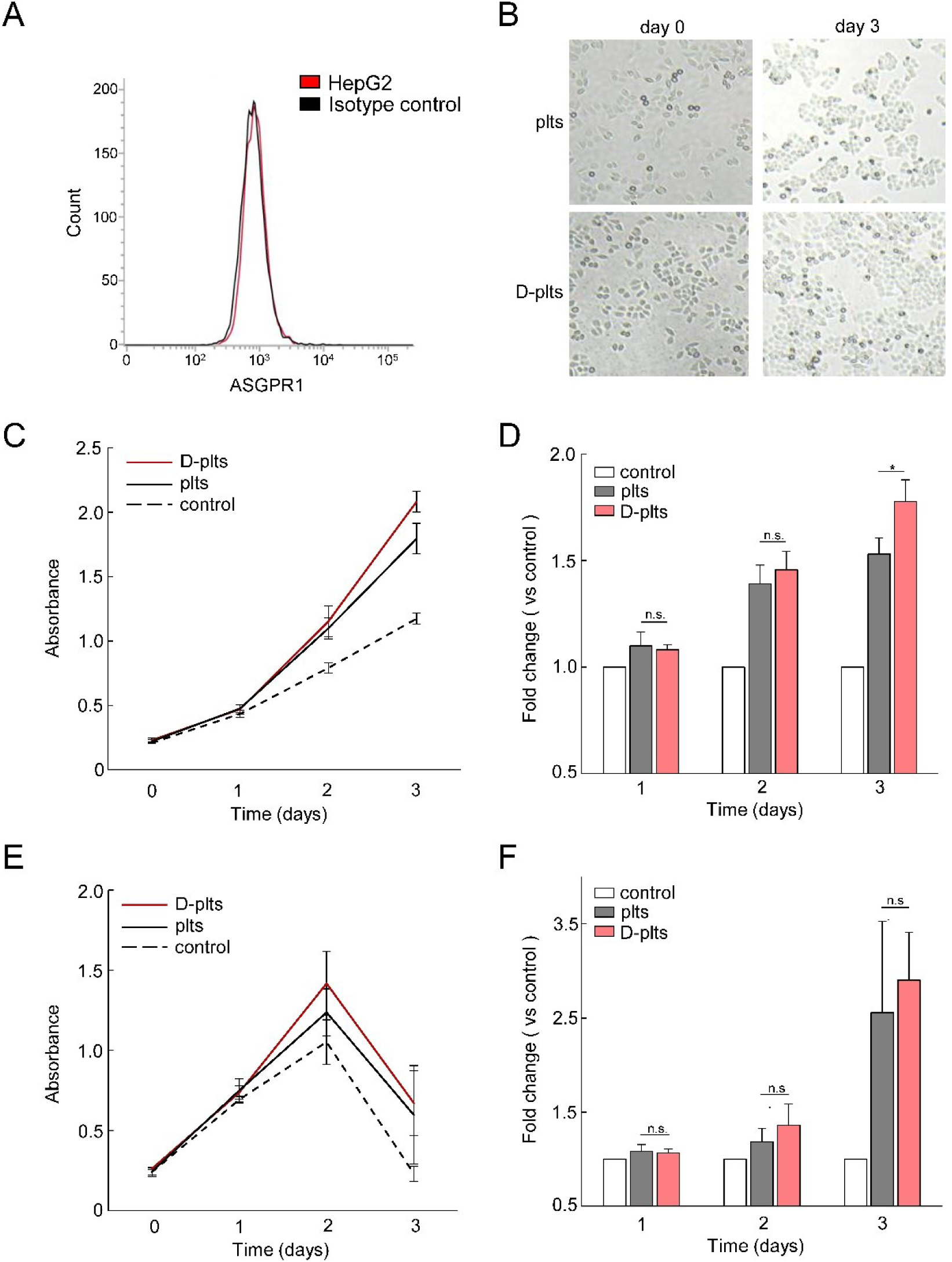
Desialylated platelets promote proliferation of HepG2 and HuH-7 cells independent of ASGPR1 expression. (**A**) ASGPR1 expression in HepG2 cells was analyzed by FCM using an anti-ASGPR1 antibody (clone 8D7). (**B**) Representative images of HepG2 cells on days 0 and 3 in the presence of D-plts or plts. (**C**) HepG2 cell proliferation was measured using a Cell Counting Kit-8 (CCK-8) assay by measuring absorbance at 450 nm (OD_450_) on days 0, 1, 2, and 3. (**D**) Fold change in absorbance was calculated by normalizing the OD_450_ value of each sample to that of the control group. (**E**) Representative images of HuH-7 cells on days 0 and 3 in the presence of D-plts or plts, as described in (**B**). (**F**) Fold changes in absorbance in HuH-7 cells were calculated as described in (**D**). Data are presented as mean ± SD (n = 3). Statistical comparisons between two groups were performed using the paired Student’s *t* test. Statistical significance was defined as \**p* < 0.05 and \*\**p* < 0.01.

To investigate whether a similar proliferative effect could be observed in other hepatocellular carcinoma cell lines, we performed analogous experiments using HuH-7 cells, which express lower levels of ASGPR1. A trend toward increased cell growth was observed in the D-plts group, although this did not reach statistical significance (Figs 2E and F).

These results suggest that D-plts promote the proliferation of hepatocellular carcinoma cell lines independently of ASGPR1 expression. The trend observed in HuH-7 cells, although not statistically significant, is consistent with this interpretation.

### Desialylated platelets promote HepG2 cell growth via ERK1/2

Because D-plts promoted HepG2 cell growth, we next investigated the intracellular signaling pathways underlying this effect. The JAK2–STAT3 and protein kinase B (Akt)/phosphatidylinositol 3-kinase (PI3K) signaling pathways are known to regulate cell proliferation and survival [17]. We examined the activation of these pathways by assessing the phosphorylation of JAK2, STAT3, and Akt using immunoblotting. Neither D-plts nor normal plts induced activation of these signaling pathways (Fig 3A). In contrast, phosphorylation of extracellular signal-regulated kinase 1/2 (ERK1/2) was moderately increased in HepG2 cells 15 min after the addition of D-plts compared with normal plts (Figs 3B and C).

**Fig 3.**
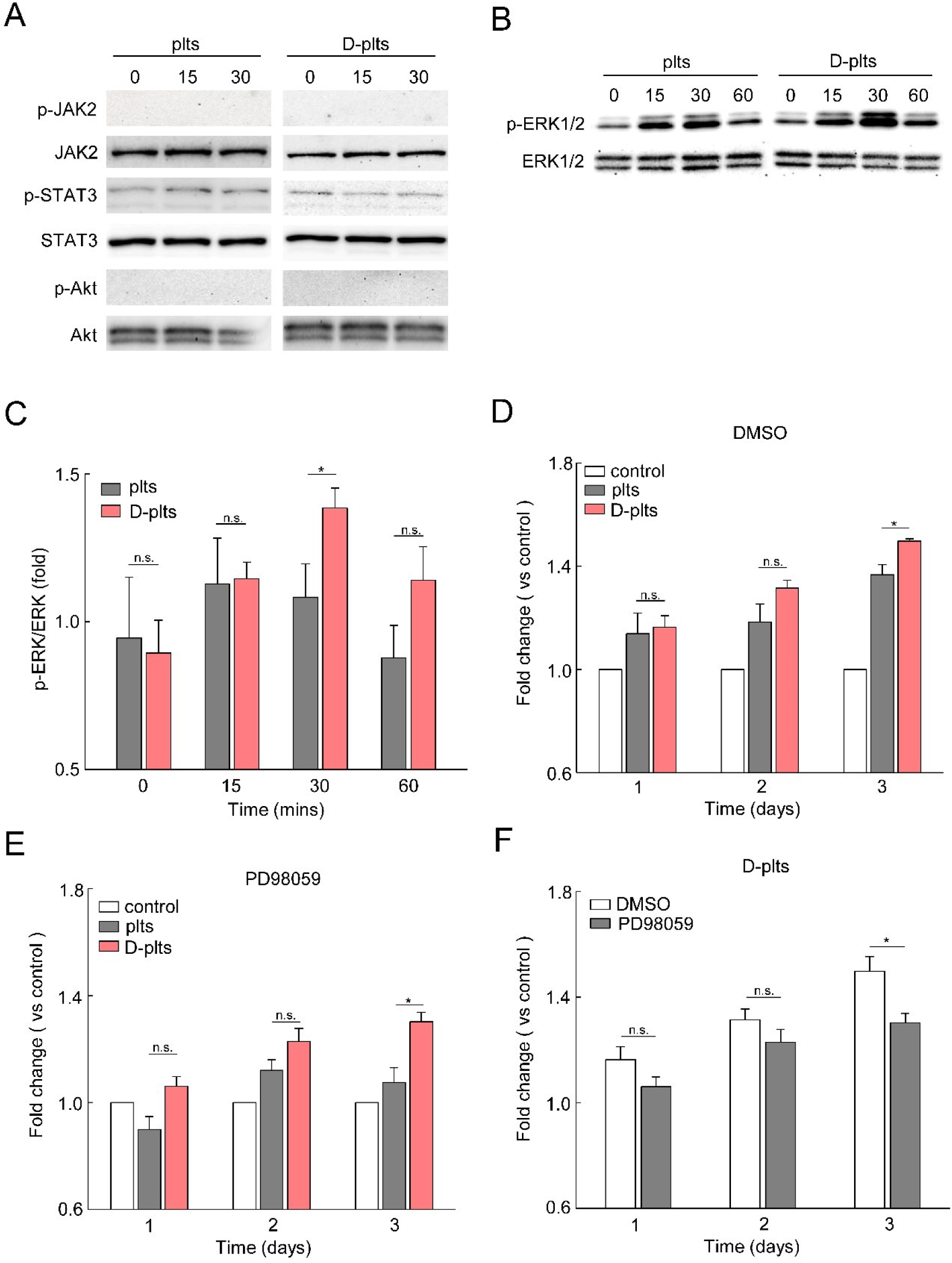
Desialylated platelets promote HepG2 cell proliferation through ERK1/2 signaling. (**A**) The levels of phospho-JAK2, total JAK2, phospho-STAT3, total STAT3, phospho-Akt, and total Akt were analyzed by immunoblotting after treatment with D-plts or plts. (**B**) The levels of phospho-ERK1/2 and total ERK1/2 were analyzed by immunoblotting. (**C**) Immunoblot bands were quantified by densitometric analysis using ImageJ and normalized to the corresponding total protein levels. (**D, E**) HepG2 cell proliferation in the presence of DMSO (**D**) or the MEK inhibitor PD98059 (10 µM) (**E**) was evaluated using a CCK-8 assay by measuring OD_450_ on days 0, 1, 2, and 3. (**F**) Comparison of D-plt-induced HepG2 cell proliferation in the presence of DMSO or PD98059 (10 μM). Data are presented as mean ± SD (n = 3). Statistical comparisons between two groups were performed using the paired Student’s *t* test. Statistical significance was defined as \**p* < 0.05 and \*\**p* < 0.01.

To further investigate the role of ERK1/2 signaling in D-plt–induced HepG2 cell growth, HepG2 cells were treated with D-plts or normal plts in the presence of dimethyl sulfoxide (DMSO) or the mitogen-activated protein kinase kinase (MEK) inhibitor PD98059, which selectively inhibits ERK1/2 activation. After 3 days of culture, D-plts significantly enhanced HepG2 cell growth compared with normal plts under DMSO conditions (Fig 3D). Although D-plts also promoted significantly greater cell growth than normal plts in the presence of PD98059 (Fig 3E), PD98059 treatment significantly reduced HepG2 cell growth compared with the corresponding DMSO-treated groups (Fig 3F). Taken together, these results suggest that ERK1/2 signaling plays an important role in the growth-promoting effects of D-plts on HepG2 cells. In contrast, no apparent activation of the canonical JAK–STAT or PI3K/Akt pathways was observed under the conditions tested. Nevertheless, the persistence of a significant growth-promoting effect in the presence of PD98059 suggests that signaling pathways other than ERK1/2 may also contribute to this response.

### Epidermal growth factor (EGF) released from desialylated platelets promotes HepG2 cell proliferation

To further evaluate the effect of indirect interactions between platelets and HepG2 cells, a transwell system was used, in which platelets were placed in the upper chamber and HepG2 cells in the lower chamber, allowing co-culture under non-contact conditions (Fig 4A). Under these conditions, both the normal plts and D-plts groups promoted HepG2 cell growth compared with the control group; however, no significant difference was observed between the normal plts and D-plts groups (Fig 4B).

**Fig 4.**
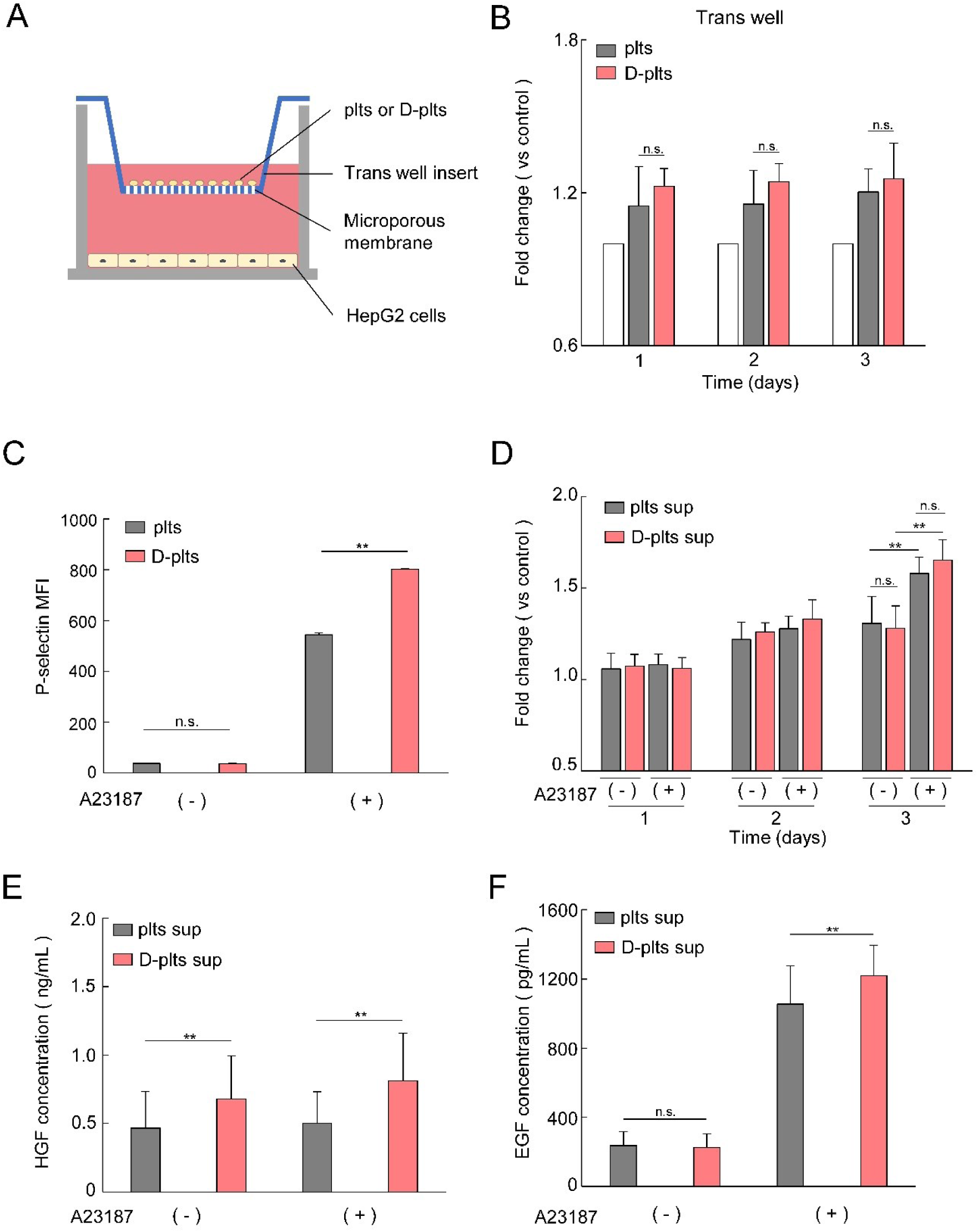
Desialylated platelets promote hepatocyte proliferation through the release of soluble factors upon activation. (**A, B**) Transwell assay. HepG2 cells were seeded in the lower chamber, whereas D-plts, plts, or HEPES (control) were added to the upper chamber. Cell proliferation was measured using a CCK-8 assay by measuring OD_450_ on days 0, 1, 2, and 3. Fold changes in absorbance were calculated by normalizing the OD_450_ value of each sample to that of the control group. (**C**) D-plts and plts were treated with the indicated concentrations of the calcium ionophore A23187. P-selectin expression was subsequently analyzed using a FITC-conjugated anti-CD62P antibody. P-selectin expression was quantified by FCM as an indicator of platelet activation. (**D**) HepG2 cell proliferation was analyzed using a CCK-8 assay following the addition of supernatants collected from A23187-stimulated D-plts or plts. Fold changes in absorbance were calculated by normalizing the OD_450_ value of each sample to that of the untreated control group. (**E, F**) Supernatants were collected after stimulation of D-plts and plts with A23187. Hepatocyte growth factor (HGF) and epidermal growth factor (EGF) concentrations in the supernatants were measured by enzyme-linked immunosorbent assay (ELISA). Data are presented as mean ± SD (n = 3). Statistical comparisons between two groups were performed using the paired Student’s *t* test. Statistical significance was defined as \**p* < 0.05 and \*\**p* < 0.01.

In contrast, under direct contact conditions, D-plts exerted a greater stimulatory effect on HepG2 cell growth than normal plts (Figs 2C and D). These findings suggest that D-plts enhance HepG2 cell growth primarily through direct cell–cell contact, whereas indirect mechanisms contribute to a lesser extent. To investigate the contribution of platelet-derived soluble factors, platelets were stimulated with A23187 (a calcium ionophore), and platelet activation was confirmed by assessing P-selectin expression. P-selectin expression was increased, particularly in D-plts (Fig 4C and S3 Fig), indicating greater platelet activation.

The supernatants from A23187-stimulated normal plts and D-plts were collected and added to HepG2 cells, resulting in a significant increase in cell proliferation in both the normal plts and D-plts groups after 3 days (Fig 4D). Subsequent analysis of the supernatants revealed that hepatocyte growth factor (HGF) levels were unchanged, whereas EGF levels were significantly higher in the supernatant from D-plts than that in that from normal plts (Figs 4E and F). In addition, EGF stimulation induced ERK phosphorylation in HepG2 cells (S4 Fig), confirming that the ERK signaling pathway is functional in these cells.

Taken together, these findings suggest that D-plts are more susceptible to activation than normal plts, as evidenced by enhanced P-selectin expression and increased EGF secretion. Moreover, platelet-derived EGF released upon activation may contribute to HepG2 cell proliferation through ERK signaling.

### Desialylated platelets promote the growth of non-malignant hCLiPs

We previously demonstrated that D-plts promote the proliferation of HepG2 cells (Figs 2B–D). To determine whether a similar effect occurs in non-malignant cells, we investigated the effect of D-plts on normal hepatocytes using hCLiPs [18–20]. hCLiPs are generated by small-molecule reprogramming of primary hepatocytes into a proliferative, liver progenitor-like state (Fig 5A), and serve as a non-malignant hepatic progenitor cell model. ASGPR1 expression was not detected in hCLiPs, as confirmed by FCM (Fig 5B).

**Fig 5.**
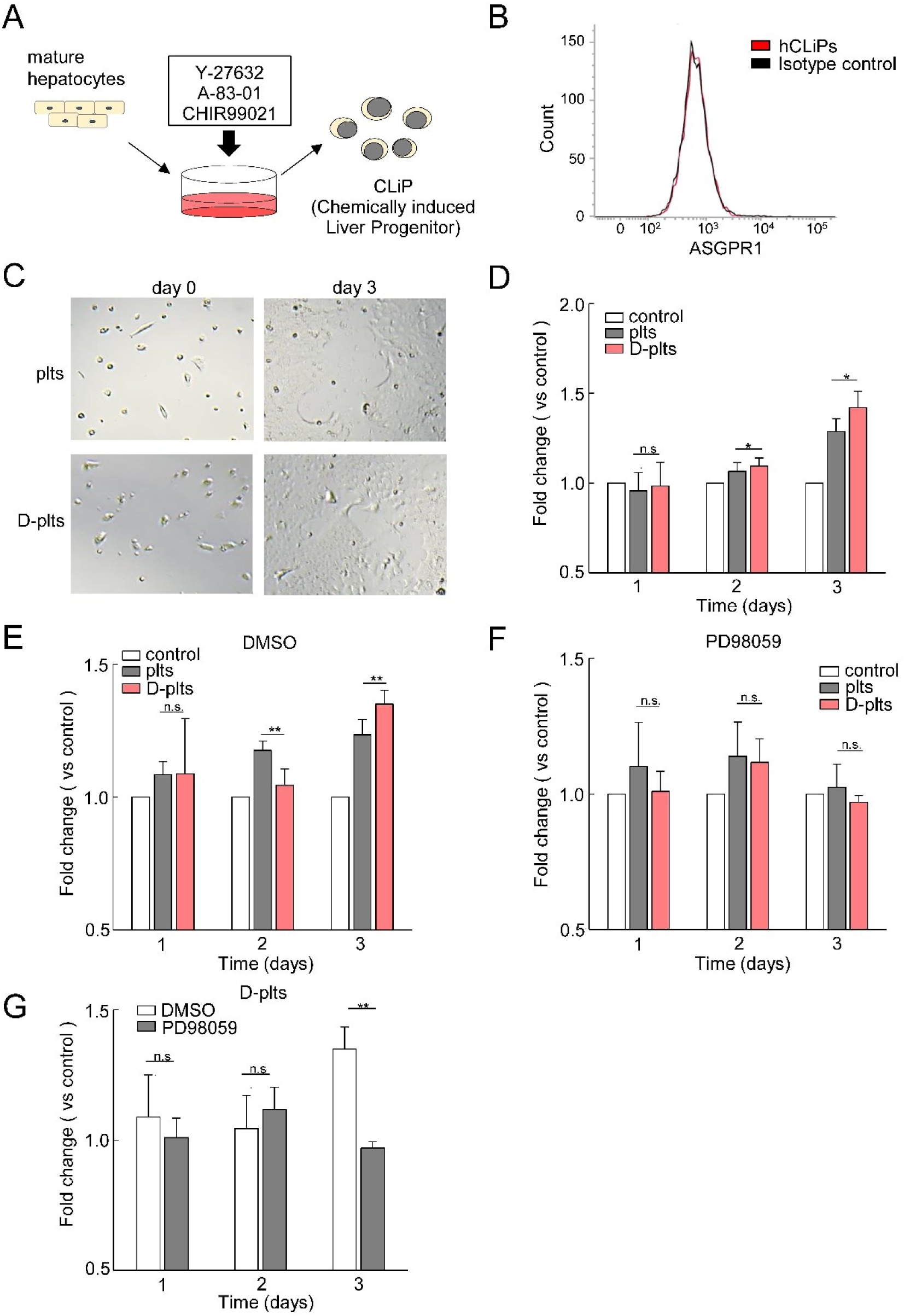
Desialylated platelets promote proliferation of hCLiPs independently of ASGPR1 expression. (**A**) Schematic representation of the induction protocol for generating hCLiPs from primary human hepatocytes using small molecules. (**B**) ASGPR1 expression in hCLiPs was analyzed by FCM using an anti-ASGPR1 antibody. (**C**) Representative images of hCLiPs on days 0 and 3 in the presence of D-plts or plts. (**D**) hCLiP proliferation was measured using a CCK-8 assay by measuring OD_450_ on days 0, 1, 2, and 3. Fold changes in absorbance were calculated by normalizing the OD_450_ value of each sample to that of the control group. (**E, F**) hCLiP proliferation in the presence of DMSO (**E**) or the MEK inhibitor PD98059 (10 µM) (**F**) was evaluated using a CCK-8 assay by measuring OD_450_ on days 0, 1, 2, and 3. (**G**) Comparison of D-plt-induced hCLiPs proliferation in the presence of DMSO or PD98059 (10 μM). Data are presented as mean ± SD (n = 3). Statistical comparisons between two groups were performed using the paired Student’s *t* test. Statistical significance was defined as \**p* < 0.05 and \*\**p* < 0.01.

When hCLiPs were co-cultured with D-plts, cell growth was significantly greater than that in the control and normal plts (Figs 5C and D). Additionally, D-plt-induced proliferation was significantly suppressed by the MEK inhibitor PD98059, suggesting that cell proliferation is mediated through the ERK signaling pathway, consistent with the mechanism identified in HepG2 cells (Figs 5E and F).

These results demonstrate that D-plts promote cell proliferation via the ERK signaling pathway not only in HepG2 cells but also in a non-malignant hepatocyte model.

## Discussion

As chronic liver disease progresses, it leads to irreversible liver dysfunction, and liver transplantation remains the only curative option [21,22]. However, donor shortages and surgical risks highlight the need for alternative therapeutic strategies. Furthermore, ASGPR expression is reduced in chronic liver disease, which may alter hepatocyte responses to circulating factors and desialylated ligands [12]. In this study, we used HepG2 and HuH-7 cells as models of hepatocytes with reduced ASGPR expression to investigate whether D-plts can promote cell proliferation under these conditions.

We demonstrated that D-plts significantly enhanced HepG2 cell proliferation compared with control and normal plts groups (Figs 2B–D). In contrast, under non-contact co-culture conditions, no difference was observed between normal plts and D-plts groups (Fig 4B), suggesting that the proliferative effect is largely dependent on direct cell–cell interactions, despite the presence of platelet-derived mitogens such as platelet-derived growth factor (PDGF), vascular endothelial growth factor (VEGF), and HGF [23–25].

Our findings differ from previous reports showing activation of the JAK2–STAT3 pathway by D-plts [10]. This discrepancy may be explained by differences in ASGPR expression. Whereas previous studies [10] suggested that D-plts activate JAK2–STAT3 signaling via ASGPR, our results indicate that D-plts primarily promote cell proliferation through the ERK1/2 pathway in cells lacking ASGPR expression. This is supported by the significant suppression of cell proliferation following ERK1/2 inhibition with PD98059 (Figs 3D–F). However, because D-plts retained a significant growth-promoting effect even in the presence of PD98059, additional signaling pathways may also contribute to the proliferative response.

The ERK pathway is commonly activated by EGF and plays a central role in cell proliferation and survival [26–29]. In the present study, D-plts released higher levels of EGF upon A23187 stimulation than with normal plts (Fig 4F). Although the transwell experiments suggested a predominantly contact-dependent mechanism, the enhanced release of EGF indicates that soluble factors also contribute to the proliferative effect. Therefore, we propose that D-plts, owing to their increased susceptibility to activation (Figs 1E and F, Fig 4C), cooperatively activate ERK signaling through both direct cell–cell interactions and the increased availability of growth factors such as EGF.

To validate these findings in a more physiologically relevant model, we employed hCLiPs. D-plts also significantly promoted the growth of hCLiPs lacking ASGPR1 expression (Figs 5B–D). Importantly, this effect was suppressed by the MEK inhibitor PD98059, suggesting that ERK signaling represents a common downstream pathway in both HepG2 cells and hCLiPs (Fig 5F). Collectively, these results indicate that D-plt-induced hepatocyte proliferation may occur through an ASGPR1-independent mechanism, suggesting the involvement of an alternative receptor or signaling pathway.

This study has several limitations. The use of HepG2 cells, HUH-7 cells, and hCLiPs may not fully recapitulate primary hepatocytes or the in vivo liver environment. In addition, experimental desialylation and variability in platelet preparations may not fully reflect physiological processes. Further studies using primary hepatocytes and in vivo models are required to elucidate additional signaling pathways, identify the receptor(s) mediating D-plt-induced ERK activation, and assess the safety and clinical applicability of D-plt-based therapeutic strategies.

## Conclusion

Our findings demonstrate a novel role for D-plts in promoting hepatocyte proliferation through ERK signaling. These results highlight the potential of platelet glycan modifications as a basis for developing novel ASGPR1-independent therapeutic strategies for liver regeneration.

## Materials and methods

### Blood and platelet collection

Fresh whole blood samples were collected from healthy volunteers after obtaining informed consent. Blood was drawn into tubes containing 3.2% sodium citrate as the anticoagulant. The samples were centrifuged to obtain platelet-rich plasma (PRP). This study was approved by the Ethical Review Board for Medical Research Involving Human Subjects at Kumamoto Health Sciences University (Approval No. 18055).

### Flow cytometry

Platelets were diluted 10-fold in PLT-HEPES (1 mM MgCl_2_, 20 mM HEPES, 136 mM NaCl, 2 mM KCl, and 0.4 mM Na_2_HPO_4_) for analysis, and each sample contained 1×10⁶ platelets. P-selectin expression was assessed as a marker of platelet activation, whereas GPIbα (CD42b) and GPIIb (CD41) expression was assessed as platelet membrane glycoproteins. P-selectin-positive platelets were detected with a fluorescein isothiocyanate (FITC)-conjugated anti-human CD62P monoclonal antibody (Becton Dickinson Inc. [BD], San Jose, CA, USA). Glycoprotein expression was assessed using phycoerythrin-cyanine 5 (PE-Cy™5)-conjugated mouse anti-human CD42b antibody (BD) and allophycocyanin (APC)-conjugated mouse anti-human CD41 antibody (BD). Surface desialylation was evaluated using FITC-conjugated RCA-I (Vector Laboratories, Burlingame, CA, USA), a galactose-specific lectin. Samples were analyzed using a flow cytometer (FACSVerse; BD Biosciences).

### Platelet aggregation test

PRP (2 × 10^5^/μL) was dispensed into glass test tubes in 180-μL aliquots for platelet aggregation analysis (Taiyo, Osaka, Japan). Samples were prewarmed for 1 min using a platelet aggregometer (PRP313M; Taiyo). Aggregation was induced by adding collagen (final concentration, 0.5 μg/mL), ADP (final concentration, 1 μM), or ristocetin (final concentration, 0.6 mg/mL) in a volume of 20 μL. Platelet aggregation was subsequently recorded.

### Preparation of desialylated platelets

Platelets were treated with 2.5 U/mL NEU from *Clostridium perfringens* (Sigma-Aldrich, St. Louis, MO, USA) at 37°C for 15 min to remove sialic acid from membrane glycoproteins. D-plts were identified by flow cytometry using RCA-I binding as described above.

### Cell culture

The human hepatocellular carcinoma cell lines HepG2 and HuH-7 were purchased from Cellular Engineering Technologies (Coralville, IA, USA) and the Japanese Collection of Research Bioresources (JCRB, Japan), respectively. Cells were cultured in Dulbecco’s modified Eagle’s medium (DMEM) (Meiji Seika Pharma Co., Ltd., Tokyo, Japan) supplemented with 10% Fetal Bovine Serum (FBS) (NICHIREI BIOSCIENCES Inc., Tokyo, Japan), penicillin G potassium, and streptomycin sulfate (Meiji Seika Pharma Co., Ltd.) at 37℃ in a humidified atmosphere containing 5% CO_2_.

hCLiPs were kindly provided by Dr. Inada [20]. The basal medium consisted of DMEM/F12 + GlutaMAX-I (Gibco, Grand Island, NY, USA) supplemented with 5 mM HEPES (Sigma, MO, USA), 30 mg/L L-proline (Sigma), 0.05% bovine serum albumin (Sigma), 10 ng/mL epidermal growth factor (Sigma), insulin-transferrin-serine-X (Life Technologies), 0.1 µM dexamethasone (Sigma), 10 mM nicotinamide (Sigma), 1 mM ascorbic acid-2 phosphate (Wako, Osaka, Japan), and an antibiotic/antimycotic solution (Life Technologies). The medium was further supplemented with 10% FBS (Life Technologies) and the small molecules 10 mM Y-27632 (Wako), 0.5 µM A-83–01 (Wako), and 3 µM CHIR99021 (Axon Medchem, Reston, VA, USA). hCLiPs were seeded onto collagen I-coated plates (IWAKI, Shizuoka, Japan) at a density of approximately 5×10^3^ viable cells/cm^2^ and cultured at 37℃ in a humidified atmosphere containing 5% CO_2_.

### Cell proliferation assay

Changes in the proliferation of HepG2 cells, HuH-7 cells, and hCLiPs were assessed using the Cell Counting Kit-8 (CCK-8; Dojindo Laboratories, Kumamoto, Japan). Cells were seeded at a density of 3 × 10^3^ cells/well and cultured for 24 h. Platelets were prepared by washing with PLT-HEPES, and D-plts were generated by NEU treatment. Platelets or D-plts were then added at a concentration of 2×10⁵ cells/well, and the cells were incubated for up to 72 h. Cell proliferation was evaluated by measuring the absorbance at 450 nm (OD₄₅₀) using a microplate reader (Multiskan FC; Thermo Scientific).

### Western blotting

Proteins were extracted from HepG2 cell lysates, separated on 12% sodium dodecyl sulfate-polyacrylamide gels, and transferred to polyvinylidene difluoride (PVDF) membranes (FUJIFILM Corporation, Tokyo, Japan). Primary antibodies against phospho-p44/42 MAPK (ERK1/2), ERK1/2, phospho-JAK2, JAK2, phospho-STAT3, STAT3, phospho-Akt, and Akt were obtained from Cell Signaling Technology (Danvers, MA, USA). Horseradish peroxidase-conjugated anti-mouse or anti-rabbit secondary antibodies (Invitrogen, Carlsbad, CA, USA) were used. Immunoblots were visualized using a chemiluminescence detection system (Bio-Rad, Hercules, CA, USA).

### Transwell co-culture assay

HepG2 cells were seeded in the lower chamber at a density of 3 × 10³ cells/well and cultured for 24 h at 37℃ in a humidified atmosphere containing 5% CO_2_. Platelets or D-plts were then added to the upper Transwell inserts (0.4-µm pore size; Corning, Corning, NY, USA), and the cells were co-cultured for 3 days. Cell proliferation was assessed using the CCK-8 assay.

### Scanning electron microscopy

Platelets were washed and treated with NEU to generate D-plts. Control platelets were treated with PLT-HEPES under identical conditions. The platelets were collected into 1.5-mL microcentrifuge tubes for subsequent analyses. Samples were fixed in 2% glutaraldehyde in 0.1 M phosphate buffer (pH 7.4) for 48 h, followed by sequential dehydration in a graded ethanol series (50%, 75%, 90%, 95%, and 100%). The ethanol was subsequently replaced with *tert*-butyl alcohol, and the samples were stored at −20°C. Frozen samples were lyophilized under vacuum, sputter-coated with platinum, and examined using a JSM-IT300 InTouchScope™ scanning electron microscope (JEOL Ltd., Tokyo, Japan) [30].

### Statistical analysis

All measurements were performed in triplicate and the results are presented as the mean ± standard deviation (SD). Comparisons between two groups were performed using a paired *t* test. Differences among multiple groups were analyzed using one-way analysis of variance (ANOVA), followed by Dunnett’s post hoc test for comparisons with the control group. Statistical significance was defined as *p* < 0.05. All statistical analyses were performed using Statcel4 software (OMS Publishing Inc., Saitama, Japan), a statistical add-in for Microsoft Excel.

## Acknowledgements

The authors thank Kano Tanabe, Ph.D. (Kumamoto Health Science University), Hirotomo Nakata, M.D., Ph.D. (Kumamoto University), for valuable discussions regarding the data analysis. The authors also thank Editage for English language editing. The authors used Google Gemini 3.0 Pro (Google LLC) to correct spelling and grammatical errors and improve English readability. The authors take full responsibility for the integrity of the content.

## Authorship contributions

IN, MU, and YK contributed to the conception and design of this study. IN, TN, MU, MO, TK, KI, YT, MM, JY, and YK acquired and analyzed the data. IN and YK interpreted the data. KI and YT provided the resources. IN wrote the draft of this article. TN, YT, MM, JY edited this article and mentored this project, and YK contributed to the revision. All authors contributed to the study and approved the manuscript as submitted.

## Data availability statement

The data that support the findings of this study are available from the first or corresponding author, [I.N. and Y.K.], upon reasonable request.

## Ethics approval statement

Approval was obtained from the Kumamoto Health Science University Ethical Review of Medical Research Involving Human Subjects (Approval No.: 18055).

## Informed consent statement

Informed consent has been obtained from all individuals included in this study.

## Supporting information

**S1 Fig.**
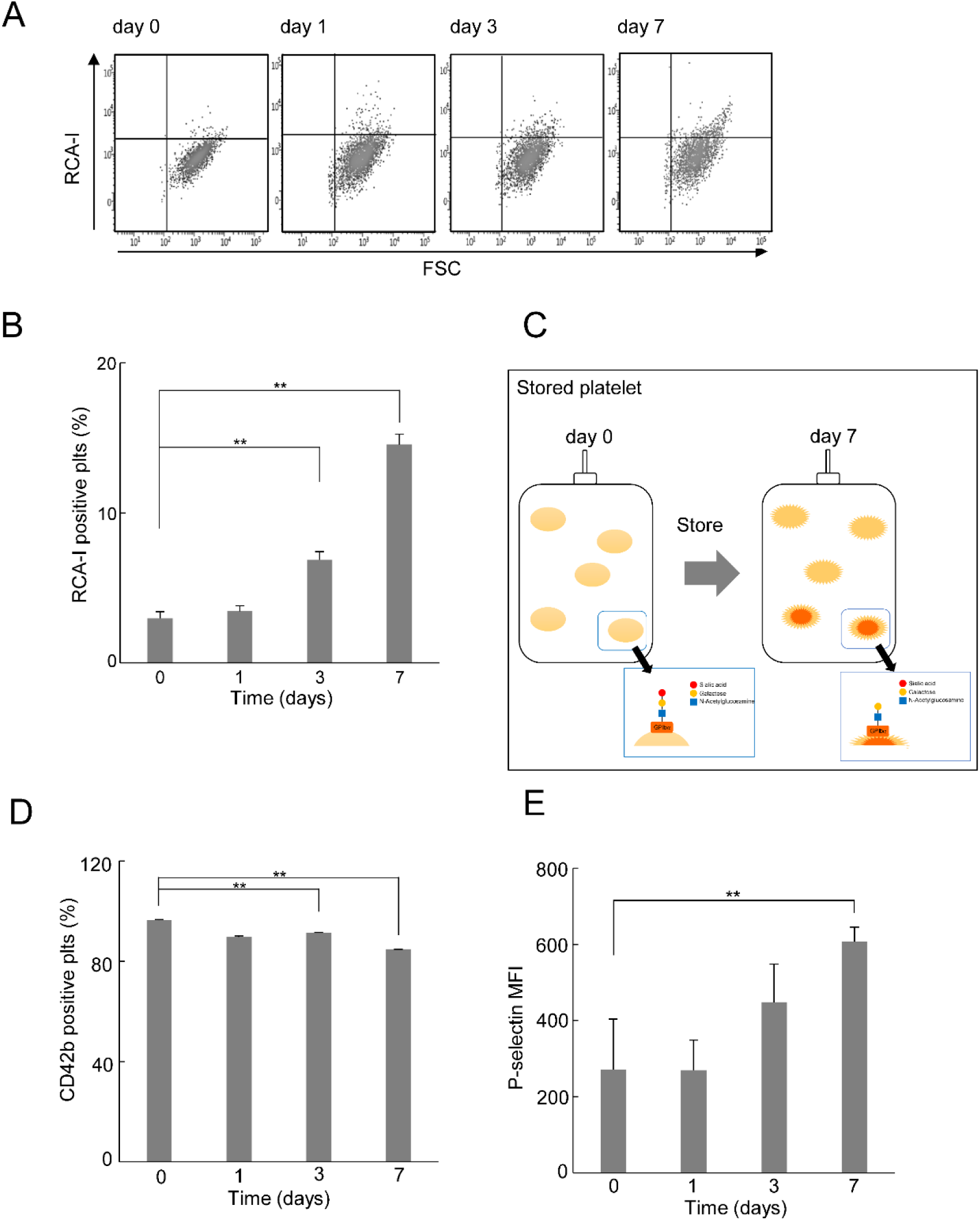
Quality and functional assessment of platelets during storage. (**A–C**) Platelet count (**A**), mean platelet volume (MPV) (**B**), and pH (**C**) were measured in platelets stored for 0, 1, 3, 7, 10, and 14 days. (**D**) Platelet aggregation was evaluated in platelets stored for 0 (black), 3 (red), 7 (green), and 14 (blue) days using collagen (final concentration: 8 μg/mL) or ADP (final concentration: 20 μM) as agonists. (**E**) Swirling was assessed by visual inspection of platelet bags under light at the indicated time points. Swirling was scored as positive (+) when clearly observed, weak (±) when faint, and negative (−) when absent. Platelet aggregation in stored bags was also evaluated visually and scored as negative (−) when no aggregates were present, weak (±) when aggregates were detectable under a microscope, and positive (+) when visible to the naked eye. (**F**, **G**) Representative SEM images of normal platelets (plts) and desialylated platelets (D-plts) under monoculture (upper panels) and co-culture with HepG2 cells (lower panels). Platelets are indicated by yellow arrows. Magnification and scale bars (μm) are shown in white. Quantitative data are presented as mean ± standard deviation (SD) (n = 3). Statistical significance was determined using Dunnett’s multiple comparison test with day 0 as the control. \**p* < 0.05, \*\**p* < 0.01.

**S2 Fig.**
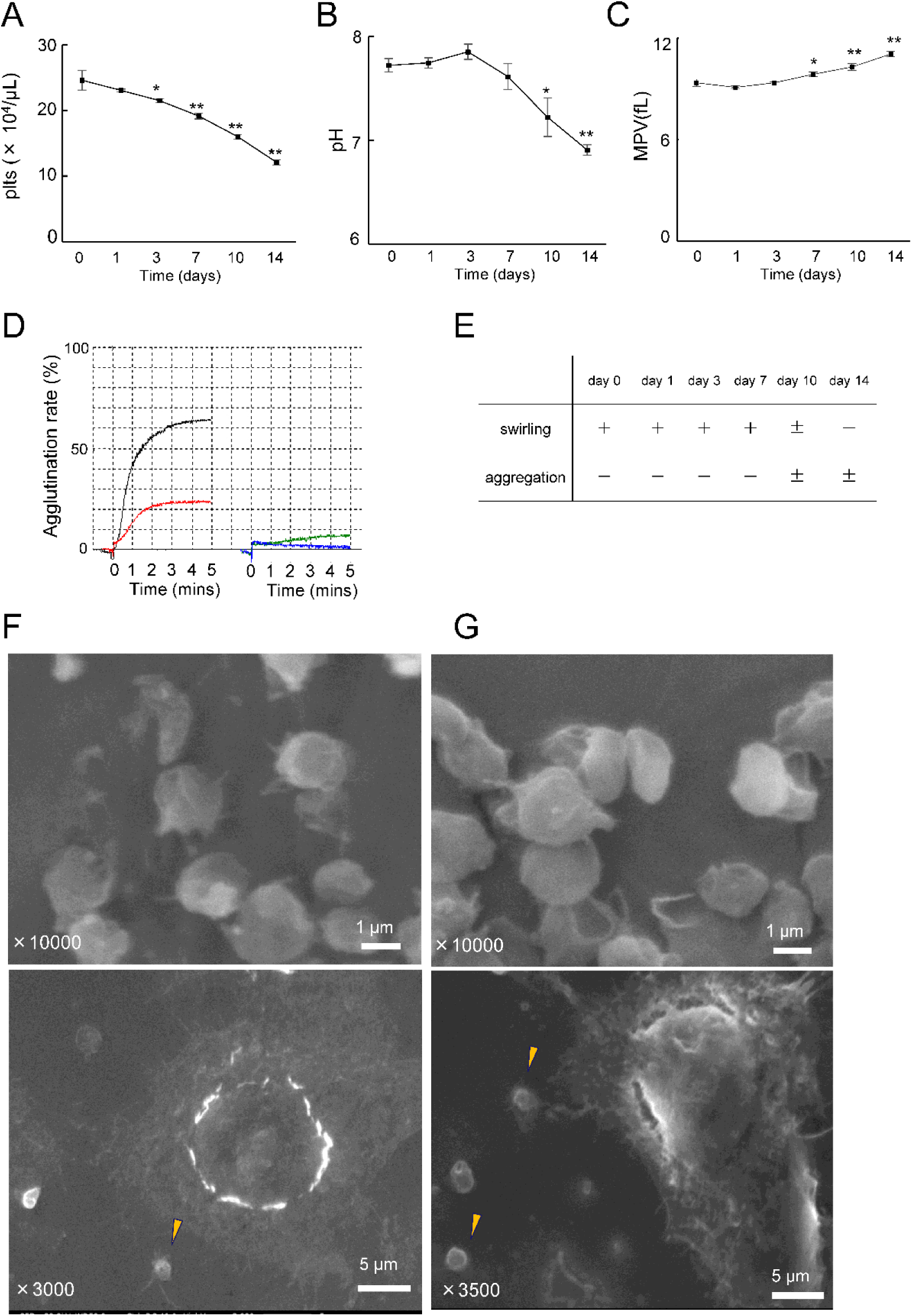
Storage-dependent changes in platelet activation and desialylation during blood bag storage. (**A, B**) Desialylation of stored platelets was evaluated by flow cytometry using FITC-conjugated *Ricinus communis* agglutinin I (RCA-I). (**A**) Representative dot plots of RCA-I staining in platelets stored in the indicated durations. (**B**) Time-dependent changes in the percentage of RCA-I–positive platelets. (**C**) Schematic illustration of changes in platelet surface glycan structures during storage. (**D**) Time-dependent changes in the percentage of CD42b-positive platelets during storage. (**E**) Time-dependent changes in the percentage of CD62P-positive platelets. Data are presented as mean ± SD (n = 3). Statistical analysis was performed using Dunnett’s multiple comparison test, with day 0 as the control. \**p* < 0.05, \*\**p* < 0.01.

**S3 Fig.**
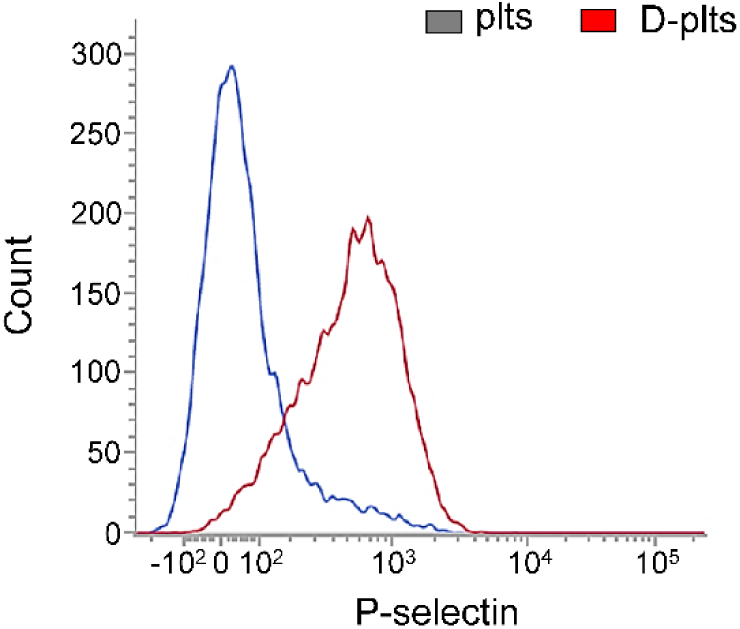
Effects of desialylation on platelet activation induced by A23187. Plts and D-plts were stimulated with A23187 (10 μM). P-selectin (CD62P) expression was evaluated by flow cytometry following staining with a FITC-conjugated anti-CD62P antibody. Representative histograms are shown.

**S4 Fig.**
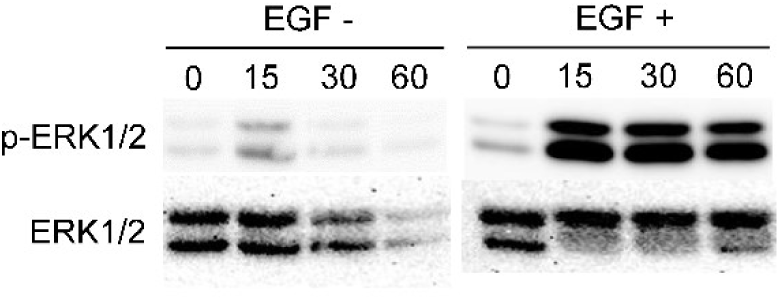
Activation of ERK signaling in HepG2 cells stimulated with recombinant EGF. HepG2 cells were stimulated with recombinant epidermal growth factor (EGF; 10 ng/mL) for 0, 15, 30, and 60 min. Cell lysates were subjected to Western blot analysis using antibodies against phosphorylated ERK (p-ERK) and total ERK. Representative immunoblots are shown.

